# Conditional stability of HY5 through the ATE N-degron pathway regulates environmental responses in *Arabidopsis thaliana*

**DOI:** 10.64898/2026.02.10.705009

**Authors:** Charlene Dambire, Isabel Manrique Gil, Jorge Vicente, Carolina Farias Saad, Andreas Bachmair, Kris Gevaert, Frank Van Breusegem, Michael J. Holdsworth

**Affiliations:** School of Biosciences, University of Nottingham, LE12 5RD, UK; Max Perutz Labs, University of Vienna, Vienna BioCenter, 1030 Vienna, Austria; Department of Biomolecular Medicine, Ghent University, Technologiepark 75, 9052, Ghent, Belgium, VIB-UGent Center for Medical Biotechnology, VIB, Technologiepark 75, 9052, Ghent, Belgium; Department of Plant Biotechnology and Bioinformatics, Ghent University, Technologiepark 79, 9052, Ghent, Belgium, VIB Center for Plant Systems Biology, Technologiepark 79, 9052, Ghent, Belgium; Department of Plant Molecular Genetics, Centro Nacional de Biotecnología CSIC, C/Darwin, 3, Campus of Cantoblanco, 28049 Madrid, Spain

## Abstract

The N-degron pathways of ubiquitin mediated proteolysis target proteins for degradation dependent on their amino terminal residue, often produced after protease activity. Very few substrates have been identified in plants even though enzymes of these pathways are highly conserved in eukaryotes. Here we identify ELONGATED HYPOCOTYL5 (HY5), an important transcriptional regulator involved in many aspects of plant development, as a target for the endopeptidase METACASPASE (MC)9, producing the carboxy-terminal protein fragment (proteoform) E59-HY5. E59-HY5 is shown to be a substrate of the arginyl transferase (ATE) N-degron pathway, and influences physiological processes known to be controlled by HY5, including photomorphogenesis and the unfolded protein response. Conditional stability of E59-HY5 was shown to result from environmentally controlled ATE function, which may highlight a general mechanism for N-degron pathway regulation of proteoform and proteome function during growth and development.

## Main

Following cleavage of proteins by endopeptidases, amino (N) and carboxy (C) terminal proteolytic fragments are created. For C-terminal protein fragments (proteoforms) this results in a novel N-terminal residue that can be targeted, dependent on the nature of the residue, by N-degron pathways of the Ubiquitin Proteasome System (UPS)^1^. Enzymes of these pathways are highly conserved in eukaryotes, and in animal systems regulate many important physiological processes^2^. In plants the only known mechanism for oxygen-sensing is through the PLANT CYSTEINE OXIDASE (PCO) N-degron pathway^3^, and other N-degron pathways are involved in the plant immune response^4,5^ and thermotolerance^6^. Very few substrates of the plant N-degron pathways have been identified, and only substrates of the PCO pathway were shown to have a pathway-associated physiological role dependent on conditional stability^7-9^. A lack of known substrates is primarily due to paucity of data on intracellular protease targets in plants, even though *A. thaliana* contains more than 600 proteases^10^. One notable exception is datasets derived from N-terminal proteomic analysis of targets of the cysteine protease METACASPASE9 (AtMC9, AtMCA-IIf) from seedlings and roots^11,12^. Here we characterise one target of MC9, ELONGATED HYPOCOTYL5 (HY5). HY5 is a bZIP transcription factor that has a multitude of roles in plant growth and development, primarily characterised as a positive regulator of light-induced development, including repression of hypocotyl elongation^13^. We show that MC9 cleavage of HY5 produces a conditionally stable C-terminal proteoform E59-HY5. This proteoform has distinct fates in HY5-regulated processes including response to light and as part of the unfolded protein response (UPR). These data suggest a general principle that conditional stability of N-degron regulated C-terminal proteoforms may play important roles in controlling plant growth and development and environmental responses.

### E59-HY5 is a physiological substrate of the ATE N-degron pathway

Analysis of published degradomes of MC9 revealed HY5 as a substrate of recombinant MC9 in an *in vitro* reaction with total extracted protein from *A. thaliana* seedlings and three week old roots^11,12^. In both datasets HY5 was observed cleaved once at the P1 site R58, that is before the bZIP DNA binding domain, after the COP1 interacting domain and adjacent to PPK1 phosphorylation sites^14,15^ (Fig. 1a). Cleavage after R58 produces a C-terminal proteoform with amino-terminal E59, a destabilising residue that is a potential substrate for arginyl-transferase (ATE) activity of the ATE N-degron pathway (Fig. 1b). Both the presence of R at the P1 site (required for MC9 cleavage) and E/D (both substrates for ATE) at the P1’ site are conserved in HY5-like sequences in angiosperms, but not in more distantly related land plants, and not in the HY5 paralogue HYH (Fig. 1c, Extended Data Fig. 1). As this sequence is outside known domains of HY5 it may suggest functional significance of the conserved residues. We first confirmed the *in vitro* cleavage of HY5 by MC9 by incubating both proteins together. We used the Ubiquitin Fusion Technique (UFT) to synthesise full length HY5 protein (all constructs used in this study Extended Data Fig. 2). Whereas cleavage of UFT-HY5-GFP-3HA produced a band of the expected size for E59-HY5-GFP-3HA, UFT-HY5(R58A)-GFP-3HA (that removes the protease cleavage site) did not (Fig. 1d) confirming the previous proteomic-derived observations of HY5 as a target of MC9^11,12^. To understand the significance of cleavage by MC9 to reveal E59-HY5, the stability of the C-terminal proteoform was analysed using the UFT (Extended Data Fig. 2)^9,16^. Initially stability of E59-HY5-GFP-3HA was assessed in rabbit reticulocyte lysate, a heterologous system that contains conserved mammalian Arg/N-degron pathway components including Ate1 and Ubr1^17^. Using this approach, E59-HY5-GFP-3HA was shown to be unstable, but stabilised in the presence of bortezomib (BZ, an inhibitor of the 26S proteasome) (Fig. 1e). Assessing the transgene *35S:UFT-E59-HY5-GFP-3HA* in *A. thaliana hy5-2* 7-day old light-grown seedlings showed that the abundance of E59-HY5-GFP-3HA is regulated by the ATE N-degron pathway (Fig. 1f, Extended Data Fig. 3). In *hy5-2* its abundance was very low but was increased in the nucleus by BZ, and by E59A substitution (Ala is not a destabilising residue). In the absence of N-degron pathway enzyme components PRT6 or ATE1 (ATE activity is encoded in *A. thaliana* by two genes, *ATE1* and *ATE2*), the stability of E59-HY5-GFP-3HA also increased. These data demonstrate that E59-HY5 is a substrate of the ATE N-degron pathway *in vivo*, and that for E59-HY5, ATE1 is the dominant regulator of degradation compared to ATE2. Although the multifunctional protein BIG has previously been shown to potentiate the activity of PRT6 in enhancing degradation of substrates of the PCO N-degron pathway^18^, for E59-HY5-GFP-3HA we did not observe a greater stabilisation in *prt6-1 big-2* than that in *prt6-1*.

**Figure 1:**
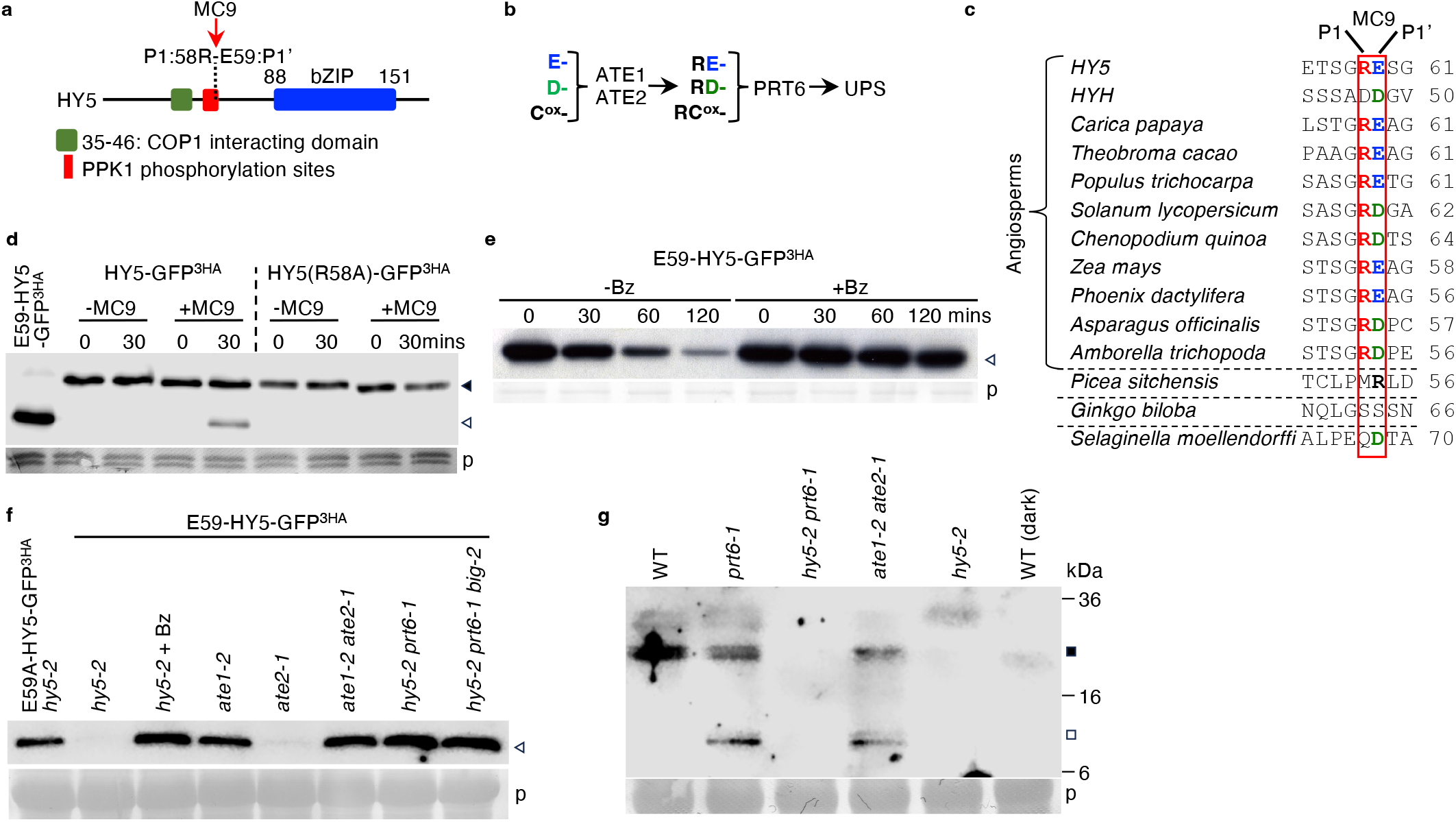
METACASPASE9 cleaves HY5 to reveal N-terminal E59. a. Schematic presentation of the HY5 protein indicating important domains and residue positions. The P1 R58 and P1’ E59 positions of the MC9 cleavage site are indicated. bZIP; DNA binding domain b. Diagrammatic representation of the ATE N-degron pathway. Single amino acid codes are used; ATE, arginyl-transferase (2 genes in *A. thaliana*); PRT6, PROTEOLYSIS6; UPS, Ubiquitin Proteasome System; ox indicates oxidised Cys residue. c. Evolutionary conservation of the P1 and P1’ residues at the MC9 recognition site in HY5 like proteins from different plant lineages. Destabilising residue colour as in b. d. *In vitro* cleavage of HY5-GFP-3HA by MC9. Comparison of time course of cleavage of HY5 with HY5(R58A) that abolishes the MC9 P1 recognition site. e. *In vitro* stability of E59-HY5-GFP-3HA over time in rabbit reticulocyte lysate in the presence or absence of bortezomib (BZ). f. *In vivo* stability of E59-HY5-GFP-3HA or E59A-HY5-GFP-3HA in *hy5-2* or N-degron pathway mutant seedlings (7 day old seedlings grown in constant light). g. Endogenous HY5 stability in WT, *prt6-1, hy5-2 prt6-1* and *hy5-2* (using polyclonal anti-HY5 antibody). Plants were grown in in the dark for 5 days and then transferred to the light for 6hours. Ponceau staining of Western blots is shown (p). Anti HA primary antibody unless otherwise stated. Closed triangle, full length HY5; Open triangle, HA-tagged HY5 Ct-proteoform; closed square, full length HY5; open square, HY5-related proteoform.

To confirm that HY5 is cleaved *in vivo* we analysed the endogenous HY5 protein by Western blot analysis during early seedling growth and establishment, where MC9 has been shown to be active^11,19^. Using an anti-HY5 antibody we observed uncleaved HY5 only in samples from seedlings exposed to light. We also observed that a protein fragment of equivalent size to E59-HY5 accumulated in 5-day old etiolated seedlings transferred to light for 6 hours in *prt6-1* and *ate1 ate2*, but not Wild Type (WT; accession Col-0), *hy5-2 prt6-1* or *hy5-2*, and only in the light (Fig. 1g). These results demonstrate that HY5 is cleaved during early seedling development to reveal a proteoform that is degraded through the ATE N-degron pathway.

### E59-HY5 plays physiological roles in HY5-mediated processes

To understand the physiological importance of the E59-HY5 C-terminal proteoform we analysed two processes in which HY5 has been shown to exert a major effect, and that involve proteasomal destruction of HY5; the transition from skotomorphogenesis to photomorphogenesis^20^, and the Unfolded Protein Response (UPR)^21^. During skotomorphogenesis HY5 is constitutively degraded through the COP1-SPA complex, but following transfer to light, extra-nuclear sequestration of COP1 allows accumulation of HY5 and activation of light responses (including inhibition of hypocotyl elongation)^22-24^. E59-HY5 does not contain the COP1-interacting domain (residues 35-46) (Fig. 1a) and therefore evades COP1 regulation. Conversely, it was shown that HY5 represses the expression of genes that activate the UPR, and HY5 is degraded by the UPS in seedlings through an unknown mechanism, to relieve repression in response to the UPR trigger tunicamycin^21^.

As previously reported hypocotyl elongation was not inhibited in the light in *hy5-2* compared to WT (Fig. 2a, Extended Data Fig. 4a,b)^25^. In contrast, *prt6-1* and *ate1-2* had shorter hypocotyls than WT, indicating that a physiological substrate of these enzymes that represses elongation is present in seedlings and normally removed through their activities. Hypocotyl length in *hy5-2 prt6-1* and *hy5-2 ate1-2 ate2-1* was much greater than *prt6-1* or *ate1-2 ate2-1* indicating that a HY5 proteoform containing a destabilising residue targeted by the N-degron pathway is stabilised to repress elongation. In *hy5-2 prt6-1* and *hy5-2 ate1-2* hypocotyl length was strongly reduced by *35S:UFT-E59-HY5-GFP-3HA* and in *hy5-2* by *35S:UFT-E59A-HY5-GFP-3HA*. These data indicate that the E59-HY5 proteoform is responsible for reduced hypocotyl length of *prt6-1*. Hypocotyl length in *35S:UFT-E59-HY5-GFP-3HA hy5-2* was also reduced in comparison to *hy5-2*, indicating that this proteoform is not completely unstable in the presence of a functioning N-degron pathway. Full length HY5 expressed through UFT also reduced hypocotyl length in a *prt6-1* dependent manner. To determine whether a *HY5-*derived proteoform could repress hypocotyl elongation in an endogenous context we analysed the influence of mutant versions of the HY5 protein in the *hy5-2* background, driven by a 756bp promoter fragment previously shown to provide HY5 functionality^26^ (Fig. 2a, Extended Data Fig.2). Whereas *promHY5:HY5-GUS hy5-2* repressed hypocotyl elongation like WT, *promHY5:HY5(E59A)-GUS* (where ATE-regulated degradation does not occur) substitution resulted in reduced hypocotyl length, whereas *promHY5:HY5(R58A)-GUS* (in which the MC9 cleavage site is removed) did not. This demonstrates the importance of the R58 cleavage site, and E59 as the N-terminal residue of the C-terminal proteoform.

**Figure 2:**
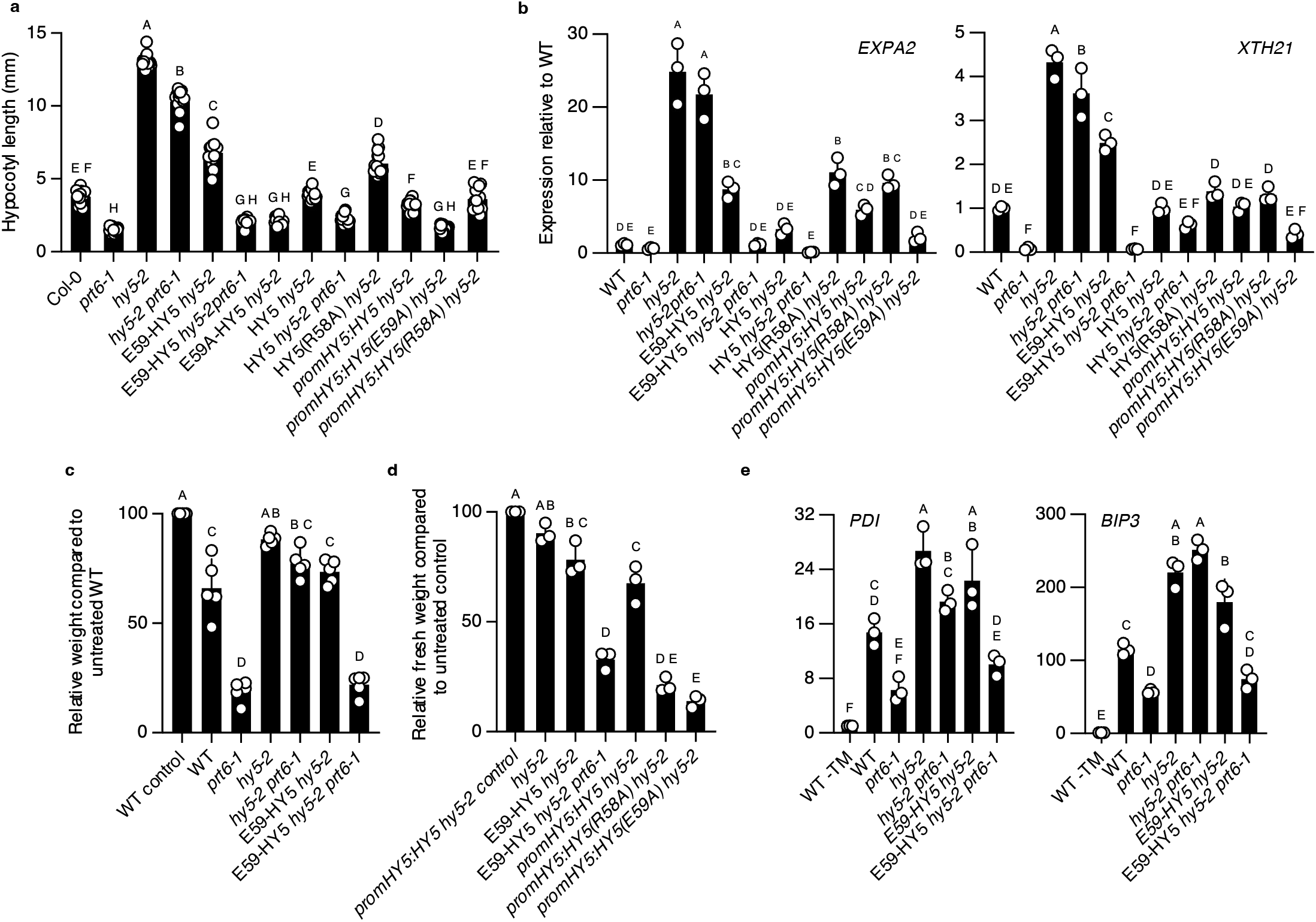
Physiological importance of the E59-HY5 C-terminal proteoform. a. Hypocotyl length of 7-day old seedlings grown in short days of WT, *hy5-2* and N-degron pathway mutant combinations. Data are presented as mean ± SD (n = 15) significant differences denoted with letters for one-way ANOVA (adjusted p<0.05). b. Expression of *HY5*-regulated genes *EXPA2* and *XTH21* in 7-day old seedlings grown in short days of WT, *hy5-2* and N-degron pathway mutant combinations. Data are presented as mean ± SD (n = 3) significant differences denoted with letters for one-way ANOVA (p<0.05). c,d. Relative weights of 14-day old light grown seedlings of WT, *hy5-2*, N-degron pathway mutant combinations and promoter constructs, seedlings grown in constant light on 60ng/ml Tunicamycin (TM). Data are presented as mean ± SD (c, n = 5; d, n = 3) significant differences denoted with letters for one-way ANOVA (adjusted p<0.05). e. Expression of *UPR*-regulated genes *BIP3* and *PDI* in 14-day old light grown seedlings grown in constant light on 60ng/ml Tunicamycin of WT, *hy5-2* and N-degron pathway mutant combinations. Comparison of expression with untreated WT seedlings. Data are presented as mean ± SD (n = 3) significant differences denoted with letters for one-way ANOVA (p<0.05).

We analysed the influence of the N-degron pathway on gene expression related to hypocotyl elongation in 7-day old seedlings grown in short days. Transcripts for cell wall modifying enzymes *XTH21* (AT2G18800) and *EXPA2* (AT5G05290) were previously shown to be repressed by *HY5* in the regulation of hypocotyl elongation^27^. Expression of these genes was repressed by E59-HY5-GFP-3HA in a *PRT6* dependent manner (Fig. 2b), indicating that E59-HY5 influences gene expression. In addition, E59-HY5-GFP-3HA repressed transcript accumulation in the *hy5-2* background, indicating that this fragment shows some stability under these growth conditions in a genetic background with a functional N-degron pathway. The *promHY:HY5(E59A):GUS* transgenic line showed a stronger repression of expression of both genes compared to *promHY:HY5:GUS* in *hy5-2*, indicating that within the context of the entire gene region E59 is an important residue.

As previously reported^21^, tunicamycin (TM) treatment of seedlings inhibited growth of WT but to a lesser extent *hy5-2* (Fig. 2c, Extended Data Fig. 5). Seedling growth was more strongly inhibited by tunicamycin in *prt6-1* and *ate1-2* compared to WT, but this inhibition was reduced in *hy5-2 prt6-1* and *hy5-2 ate1-2* indicating that repressed growth of this N-degron mutant was caused by *HY5*. Expression of E59-HY5-GFP-3HA in *hy5-2 prt6-1* led to a phenotype strongly resembling that of *prt6-1*. In *hy5-2, promHY5:HY5-GUS* restored a WT-like phenotype of reduced growth on tunicamycin, and both *promHY5:HY5(E59A)-GUS* and *promHY5:HY5(R58A)-GUS* both strongly repressed growth (Fig. 2d). These data indicate that both cleavage of HY5 at the MC9 site and proteasomal degradation of the C-terminal proteoform are important for activation of the UPR. The expression of UPR-activated genes *PDI* (AT5G38900) and *BIP3* (AT1G09080) was repressed by *HY5* during normal development and activated in response to tunicamycin treatment (Fig. 2e). Activation was enhanced in *hy5-2* and *hy5-2 prt6-1*, but repressed in *prt6-1*, indicating that the N-degron pathway activates UPR gene expression through removal of HY5. Unlike hypocotyl elongation E59-HY5-GFP-3HA was only able to restore response to tunicamycin in the *prt6-1* background, indicating that this fragment is not stable during the UPR. These results show that the E59-HY5 proteoform, regulated through the N-degron pathway has important roles in developmental and cellular response to environmental change.

### Different roles of the E59-HY5 proteoform

We investigated the half-life of E59-HY5 to determine the physiological relevance of MC9 cleavage and conditionality of E59-HY5 degradation, as we observed activity of this proteoform in repressing hypocotyl elongation (compare UFT-E59-HY5 *hy5-2* to *hy5-2* Fig. 2a). Analysis of UFT-E59-HY5-GFP-3HA showed that whereas this proteoform was stable in the dark in *hy5-2 prt6-1*, it was not in *hy5-2* (Fig. 3a). This shows that degradation of this fragment is independent of COP1, but dependent on PRT6. Following transfer to light the fragment transiently increased in abundance between 2 and 4h of illumination and was not visible at 6h. A HY5-derived fragment was also observed in WT seedlings following transfer to light (Fig. 3b). To investigate if conditional stability of the E59-HY5 proteoform is a property of HY5 or the N-degron pathway, we substituted E59 with R, a destabilising residue that converts the proteoform from a substrate of ATE to a substrate of downstream PRT6 (Fig. 1a) in the construct UFT-E59R-HY5-GFP-3HA (Extended Data Fig. 2). Whilst E59-HY5-GFP-3HA accumulated following transfer to the light E59R-HY5-GFP-3HA did not though it was stabilised in the presence of bortezomib (Fig. 3c, Extended Data Fig. 6a). Similarly, the artificial substrate E-3HA-GUS was stabilised whereas R-3HA-GUS was not (Extended Data Fig. 6b), indicating that N-terminal Glu is a signal for conditional stabilisation. The abundance of HY5-GFP-3HA increased in response to light, as did a Ct-proteoform fragment of a similar size to E59-HY5-GFP-3HA, however, this fragment was not visible in HY5(R58A), where the MC9 cleavage site is removed (Fig. 3d). As expected, *COP1* transcript declined after illumination^24^, *PRT6* did not change, *ATE1* increased slightly and *MC9* very strongly^11^ (Fig. 3e). In response to light the expression of both *EXPA2* and *XTH21* decreased in *E59-HY5-GFP-3HA hy5-2* but not in *E59R-HY5-GFP-3HA hy5-2* showing that transient stabilisation through N-terminal E59 influences gene expression (Fig. 3f, Extended Data Fig. 7a).

**Figure 3:**
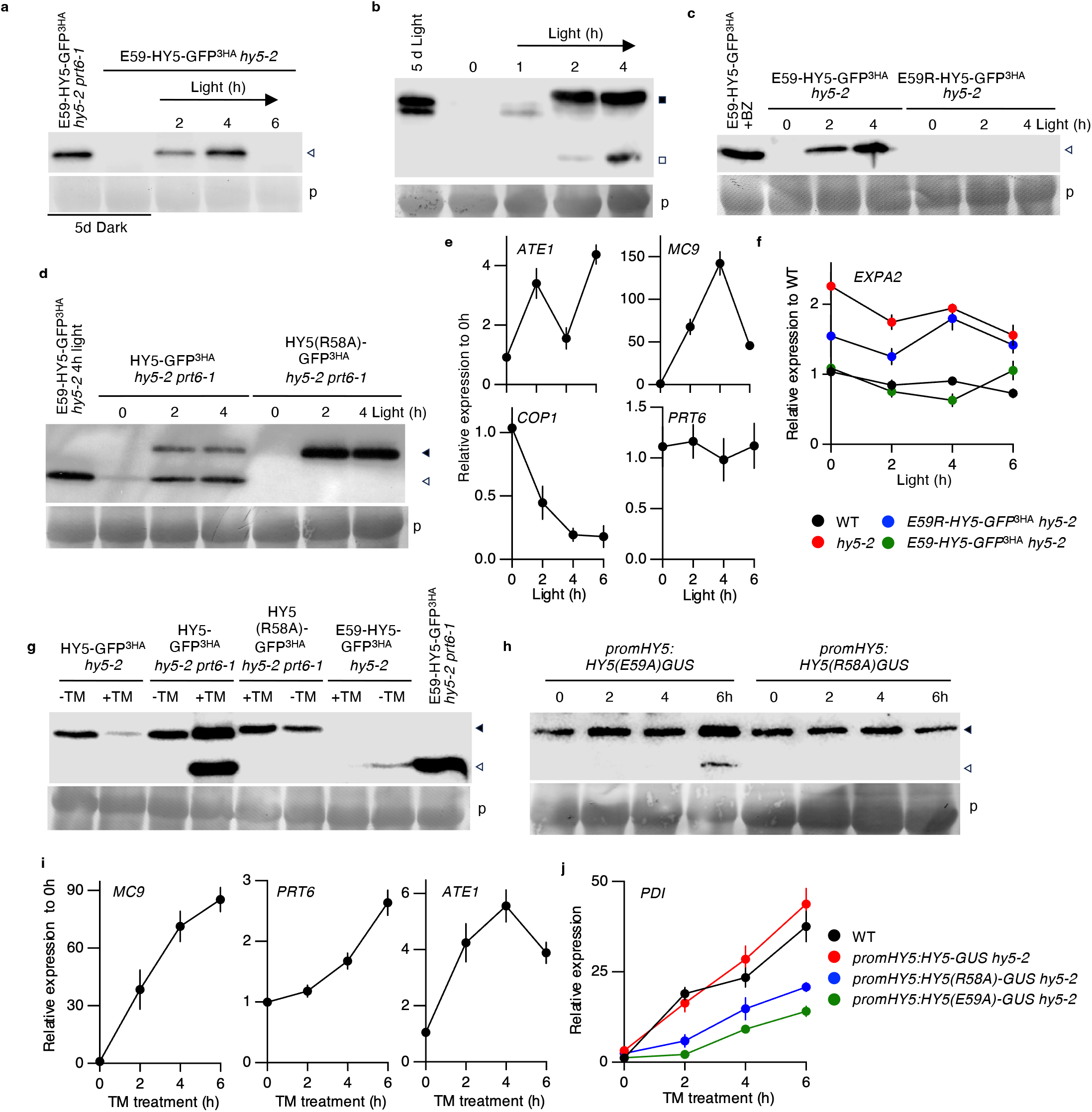
Appearance and conditional stability of E59-HY5 during seedling development. **a**. Stability of E59-HY5-GFP-3HA in *hy5-2* and *hy5-2 prt6-1* mutants in 5-day old etiolated seedlings following transfer from dark to constant light. **b**. Endogenous HY5 stability in WT. Plants were grown in in the dark for 5 days and then transferred to light for 4 hours (polyclonal anti-HY5 antibody). **c**. Stability of E59-HY5-GFP-3HA (*hy5-2*) compared to E59R-HY5-GFP-3HA (*hy5-2*) in 5-day old seedlings following transfer from dark to light. **d**. Stability of HY5-related proteoforms in 5-day old etiolated seedlings transferred to light . **e**. Expression in 5-day old etiolated seedlings of *COP1, PRT6, MC9, ATE1* following transfer from dark to light. Data are presented as mean ± SD (n = 3). **f**. Expression of *EXPA2* in WT, *hy5-2, promHY5:HY5-GUS* and *promHY5:HY5(R58A)-GUS*. Data are presented as mean ± SD (n = 3). **g**,**h**. Stability of HY5-related proteoforms in 14-day old seedlings grown on 60 ng/ml tunicamycin. Anti HA (**g**) or GUS (**h**) primary antibody. **i**. Expression in 7-day old seedlings grown in constant light of *PRT6, MC9, ATE1* during liquid incubation in tunicamycin (5 μg/ml). Data are presented as mean ± SD (n = 3). **j**. Expression of *BIP3* in WT, *hy5-2, promHY5:HY5-GUS* and *promHY5:HY5(R58A)-GUS* during liquid incubation in 7-day old seedlings grown in constant light with tunicamycin (5 μg/ml). Data are presented as mean ± SD (n = 3). Ponceau staining of Western blots is shown (p). Anti-HA primary antibody unless otherwise stated. Closed triangle, full length HY5; Open triangle, HA-tagged HY5 Ct-proteoform; closed square, full length HY5; open square, HY5-related proteoform.

As observed previously for HY5^21^, in response to growth for 14 days on tunicamycin the abundance of HY5-GFP-3HA was reduced compared to untreated seedlings (Fig. 3g). In the absence of *PRT6* a cleavage product was observed at a similar size to E59-HY5-GFP-3HA and an increase in full length HY5. In HY5(R58A) no cleavage product was observed. A similar observation was made with HY5 promoter constructs in *hy5-*2 (Extended Data Fig. 5c). Analysis of GUS accumulation in seedlings in response to growth on TM showed that either inability to cleave (R58A) or degrade (E59A) (Fig. 3h) led to relatively enhanced staining compared to the WT construct (Extended Data Fig. 8). In response to tunicamycin treatment in solution, *MC9* expression greatly increased, and to a lesser extent that of *PRT6* and *ATE1* (Fig. 3i). Expression of UPR-activated genes *BIP3* and *PDI* increased strongly in response to tunicamycin treatment in WT, and also in *hy5-2* containing the *promHY5:HY5-GUS* transgene. In comparison expression was lower in both *promHY5:HY5(R58A)-GUS* and *promHY5:HY5(E59A)-GUS* (Fig. 3j, Extended Data Fig. 7b). This indicates that either failure to cleave HY5 at R58, or persistence of the E59A proteoform both repress the UPR gene expression response.

In this work we investigated three principles of N-degron pathway biology. Firstly, we confirmed that HY5 is cleaved by a protease, MC9, to produce a C-terminal proteoform E59-HY5. We showed that this fragment is a substrate of the ATE N-degron pathway, demonstrating that substrates are introduced into the pathways following protease cleavage of target proteins. Secondly, we discovered that the stability of this proteoform transiently increased in the dark to light transition, demonstrating that conditional substrate stability can occur. Thirdly, we show that stabilisation of E59-HY5 influences hypocotyl elongation, demonstrating that conditional stability of substrates results in novel proteoform functions. In this way the N-degron pathway control of E59-HY5 stability is similar to the regulation of ERFVII stability as part of the PCO N-degron pathway, that also exhibits these three principles^9^. The discovery of this feature outside the PCO pathway suggests that conditional stability may be a general principle of N-degron pathway biology in regulating proteoform function. Previous work showed that an artificial C-terminal deletion derivative of HY5 (HY5-ΔN77, initiating at position 77 leaving just the bZIP DNA binding domain) was dissociated from COP1-regulated degradation and exhibited increased activation of photomorphogenic development compared to full-length HY5^22^. In agreement with that observation, we showed that the naturally derived E59-HY5 is also independent from COP1 regulation. Other work also demonstrated that the kinetics of COP1 migration from the nucleus in response to light occurs with a half-life of 2.4 hours, whereas full length HY5 accumulated in the nucleus by 4 hours^24^. A version of a rice HY5-like protein (OsbZIP1.2) was also shown not to contain the COP1 interacting domain as a consequence of alternative splicing^28^, and an alternatively spliced variant of HYH (altHYH) was hypothesised to enhance light-regulated gene expression as a consequence of being uncoupled from COP1 repression^29^. It is possible therefore that E59-HY5 functions with other COP1-independent factors to ensure efficient conversion to photomorphogenic development by escaping COP1-mediated degradation during the early stage of light transition. In contrast, in the UPR, E59-HY5 appears not to be functional and cleavage at R58 is a mechanism to process HY5 for proteasomal degradation.

It will be interesting to discover whether other branches of the N-degron pathways also show these three principles of substrate birth and death, which will depend on substrates being identified for each branch. Only two other plant pathway substrates have been identified outside the PCO N-degron pathway, BIG BROTHER (Y61-BB, a substrate of the E3 ligase PRT1^30^) and ATG8a (PRT7 substrate R13-ATG8a^6^), however no conditional stabilisation was investigated in either case. We used the MC9 N-terminome to uncover and investigate E59-HY5 as a potential N-degron pathway substrate, and many other potential substrates exist within this dataset^11^. It is likely therefore that many novel N-degron pathway substrates will be uncovered influencing aspects of plant growth, development and environmental responses.

## Methods

### Plant materials and growth conditions

All work was carried out in the *A. thaliana* Col-0 accession background. Mutant combinations were generated using *hy5-2* (SALK_056405)^31^, obtained from NASC (Nottingham, UK). Plants were grown as previously described^32^ unless otherwise stated.

### Generation of ubiquitin fusion technique constructs

HY5 cDNA starting at E59^11^ was PCR amplified from *Arabidopsis thaliana* and cloned into a FLAG-DHFR-UBIQUITIN-GFP-3HA vector as previously described^9^. As described in^33^ this provides a SacII restriction site downstream of the UBIQUITIN moiety that allowed cloning in frame of test proteins with defined amino-terminal residues. In this case we cloned HY5 starting at E59, so that upon cleavage *in vivo* by constitutive deubiquitinating enzymes E59-HY5GFP-3HA or equivalent proteoforms (Extended data Fig. 2) are released from the pre-protein. This construct was subcloned into plant transformation gateway vector pH7wg2^34^ providing constitutive expression from the 35S CaMV promoter. Plants were transformed as previously described^35^.

### Generation of Arabidopsis promoterHY5:GUS reporter lines

The native HY5 gene region (starting 756 nucleotides upstream of the start codon) was amplified by PCR from Col-0 genomic DNA and introduced into a Gateway compatible entry vector (PDONRzeo). The verified entry clone was then inserted into the binary destination vector pGWB3, which carries the β-glucuronidase (GUS) reporter gene at the 3’ end of the inserted DNA. The resulting construct, promHY5:HY5-GUS, was introduced into *Agrobacterium tumefaciens* strain GV3101 and used for transformation of *A. thaliana hy5-2* and *hy5-2 prt6-1* mutants as above.

### Site-directed mutagenesis

Site-directed mutagenesis primers were designed and reactions performed using the QuikChange™ protocol (Agilent-technologies). Positive clones were identified by colony PCR and confirmed by DNA sequencing. All primers used are listed in Extended Data Table 1.

### Dark/light treatments

As previously described^36^, surface-sterilized seeds were plated on ½ strength MS and chilled for 3 days at 4 °C before being transferred to continuous white light at 20 °C for 8 h to activate germination. Subsequently, unless indicated otherwise, plates were incubated in darkness at 20 °C for 5 days, then exposed to continuous light at 20 °C.

### Tunicamycin treatment

For the analysis of ER stress tolerance on plant growth, seedlings were sown on ½ strength MS supplemented with 60ng/mL Tunicamycin dissolved in DMSO or DMSO alone. The plates were chilled (4 °C) in the dark for 3days. Plates were then transferred to a growth chamber under continuous light at 20 ºC. After 14 days, plates were scanned using the EPSON PERFECTION 4990 photo scanner. Relative fresh weight of plants was measured by gently blotting seedlings on paper towel to remove surface moisture, then promptly weighing them on an analytical balance. Fresh weights of WT seedlings grown in untreated conditions were set at 100%^21^. For the time-course treatment in liquid culture, seedlings were germinated on ½ strength MS medium, chilled, and then transferred to a growth chamber under continuous light for 7 days. Following this, 30 seedlings were placed in a 6-well plate containing 3 ml of sterile water containing 5 µg/ml Tunicamycin.

### Hypocotyl length analysis

*A. thaliana* seeds were plated on ½ strength MS followed by chilling at 4ºC in the dark for 3 days, then transferred to a growth room with 8 h light, 16 h dark at 20ºC. Agar plates were placed vertically in the growth chambers. After 7 days, 15 seedlings per line were imaged for hypocotyl length using the Leica S9D Stereomicroscope. Hypocotyl length was measured using ImageJ software (http://imagej.nih.gov/ij/).

### *In vitro* protein stability analysis

The stability of proteins was assessed *in vitro* using the rabbit reticulocyte lysate coupled transcription/translation system as previously described. This heterologous system has previously been shown to contain all components of the mammalian Arg/N-degron pathway^7,37^.

### Bortezomib treatment of seedlings

Bortezomib (dissolved in DMSO) treatment was carried out in 6-well tissue culture plates (Fisher Scientific) as previously described^36^. In a total reaction volume of 3 ml, Bortezomib (Santa Cruz Biotechnology) was added to a final concentration of 25 μM. Negative controls contained an equivalent volume of DMSO (Sigma-Aldrich). 7-day-old seedlings grown in continuous light were transferred by laying the seedlings on top of the solution in each well ensuring that roots of seedlings were immersed in the reaction solution. The plate was placed on a flatbed shaker (60 rpm) for 2 h, after which material was dried with paper towel to remove excess liquid and frozen immediately in liquid nitrogen.

### Western blot analysis

*A. thaliana* seedling proteins were extracted as previously described^9^. Proteins were separated by SDS-PAGE electrophoresis and analysed by Western blotting as previously described^9^. Primary antibodies used were mouse anti-HA (Sigma-Aldrich; 1:1,000), anti-HY5 (Agrisera; 1:2,000) and anti-GUS (Sigma-Aldrich; 1:2,000). Horseradish peroxidase conjugated secondary antibodies, goat anti-Rabbit IgG (Agrisera) and goat anti-mouse IgG1 (Thermo Fisher Scientific), were used at 1:10,000 dilution. Signals were detected using LICOR WesternSure® substrate and imaged with a LICOR ODYSSEY FC Imager. Membranes were subsequently stained with Ponceau S, washed, dried, and scanned.

### RNA expression analyses

Total RNA was extracted from seedlings (age indicated for each experiment) using a RNAeasy Plant Mini Kit (QIAGEN) as previously described^9^.

### Phylogenetic analysis

Comparison of HY5 like species was carried out following BLASTp search for HY5-like sequences at NCBI. Sequence comparison was carried out using MUSCLE in MacVector v18.

### Production of recombinant MC9 protein

MC9 open reading frame was cloned into pDEST17 vector from A. *thaliana* seedling cDNA, resulting in an N-terminal fusion with a His x 6 epitope tag. Protein expression was induced at 30ºC overnight with 1mM IPTG and purified as described previously^38^.

### MC9 activity assay *in vitro*

To investigate the direct proteolytic effect of MC9 on HY5, an *in vitro* cleavage assay was performed using recombinant MC9 and HY5 substrates produced by coupled transcription/translation. HY5 WT and HY5(R58A) in the vector UFT-GFP-3HA were used as templates for *in vitro* transcription and translation (TnT® T7/T3 Coupled Reticulocyte Lysate System, Promega) according to the manufacturer’s instructions. Reactions were gently mixed on ice and incubated at 30 °C for 30 min. After incubation, 2 μl of 2.6mM cycloheximide (Sigma-Aldrich) was added to inhibit translation. Each reaction was divided into two 15 μl aliquots, one for the MC9 treatment (+MC9) and one as a negative control (−MC9). Recombinant MC9 was pre-activated prior to use in activity buffer (50 mM MES, pH 5.5, 150 mM NaCl, 10% (w/v) sucrose, 0.1% (w/v) CHAPS)^38^. For pre-activation, 20 μl activity buffer and 10 μl 100 mM DTT were added to 50 μl of recombinant MC9 (2.938 mg/mL, ∼ 63.9 µM), and the mixture was incubated at 30 °C for 10 min. For cleavage assays, 30 μl of pre-activated MC9 was added to 15 μl of the *in vitro* translation solution, followed by 10 μl activity buffer. A 25 μl aliquot was collected immediately (time 0 min). Negative control reactions contained 40 μl activity buffer in place of MC9. Reactions were incubated at 30ºC, and additional samples were subsequently collected. Reaction products were denatured by addition of 7 μl Laemmli sample buffer and water to a final volume of 42 μl, followed by incubation at 95 °C for 10 min, chilling on ice for 5 min, brief centrifugation, and storage at −20 °C.

### Green fluorescent protein imaging

7-day old *A. thaliana* seedlings grown at 20ºC in continuous light were imaged for GFP protein analysis using a Leica TCS SP8 confocal microscope (Leica Microsystems) at an excitation wavelength of 514 nm and an emission wavelength of 500 to 530 nm.

### Histochemical staining

For histochemical analysis of β-Glucuronidase (GUS) enzyme activity, plant material was incubated at 37 °C for 8 h in the dark suspended in GUS staining solution (100μM potassium ferricyanide, 100μM potassium ferrocyanide, 100 mM PBS pH 7.0, 2 mM X-gluc, 0.1% v:v Triton X-100. Seedlings were de-stained in clearing fixative (3:1 ethanol: acetic acid, 1% v:v Tween®) overnight, before mounting in Hoyer’s solution (30 g gum Arabic, 200 g chloral hydrate, 20 g glycerol, 50 ml water). Seedlings were imaged using the Leica S9D Stereomicroscope.

## Data availability

Data and resources will be shared upon reasonable request to Michael Holdsworth.

## Acknowledgements

We thank Bhavani Sundaram and Kamal Swarup for technical support of this work.

## Author information

M.J.H, C.D conceived the research with inputs from J.V, F.V.B, K.G and A.B. All authors contributed towards design of experiments and interpretation of data; M.J.H, C.,D, I.M.G., J.V, C.F.S carried out the research; M.J.H and C.D wrote the manuscript. M.J.H agrees to serve as the author responsible for contact and ensures communication. All authors read, contributed to editing and approved the manuscript.

## List of Extended Data

Extended Data Figure 1: Conservation of MC9 cleavage site in HY5 related sequences.

Extended Data Figure 2: Diagrammatical representation of constructs used in this study.

Extended Data Figure 3: Subcellular localization of E59-HY5-GFP-3xHA construct in different genetic backgrounds.

Extended Data Figure 4: Influence of ATE1 and ATE2 on HY5 regulated hypocotyl elongation, and examples of phenotypes of seedling response to light.

Extended Data Figure 5: Influence of N-degron pathway mutants on HY5 regulated response to Tunicamycin.

Extended Data Figure 6: Western blot analysis of HY5 and GUS constructs.

Extended Data Figure 7 | Expression of HY5 regulated genes in seedlings.

Extended Data Figure 8 | Histochemical analysis of promHY5:HY5-GUS constructs in seedlings.

Extended Data Table 1: List of oligonucleotide primers used in this study.

