## Supplementary Figures for "Conditional stability of HY5 through the ATE N-degron pathway regulates environmental responses in *Arabidopsis thaliana*"

**Extended Data Figure 1:** Conservation of MC9 cleavage site in HY5 related sequences.
The COP1 interacting domain (green box), MC9 cleavage site (red box ) and bZIP DNA binding domain (blue box) are shown.
Shading indicates conserved residues; dark = conserved, light = similar.

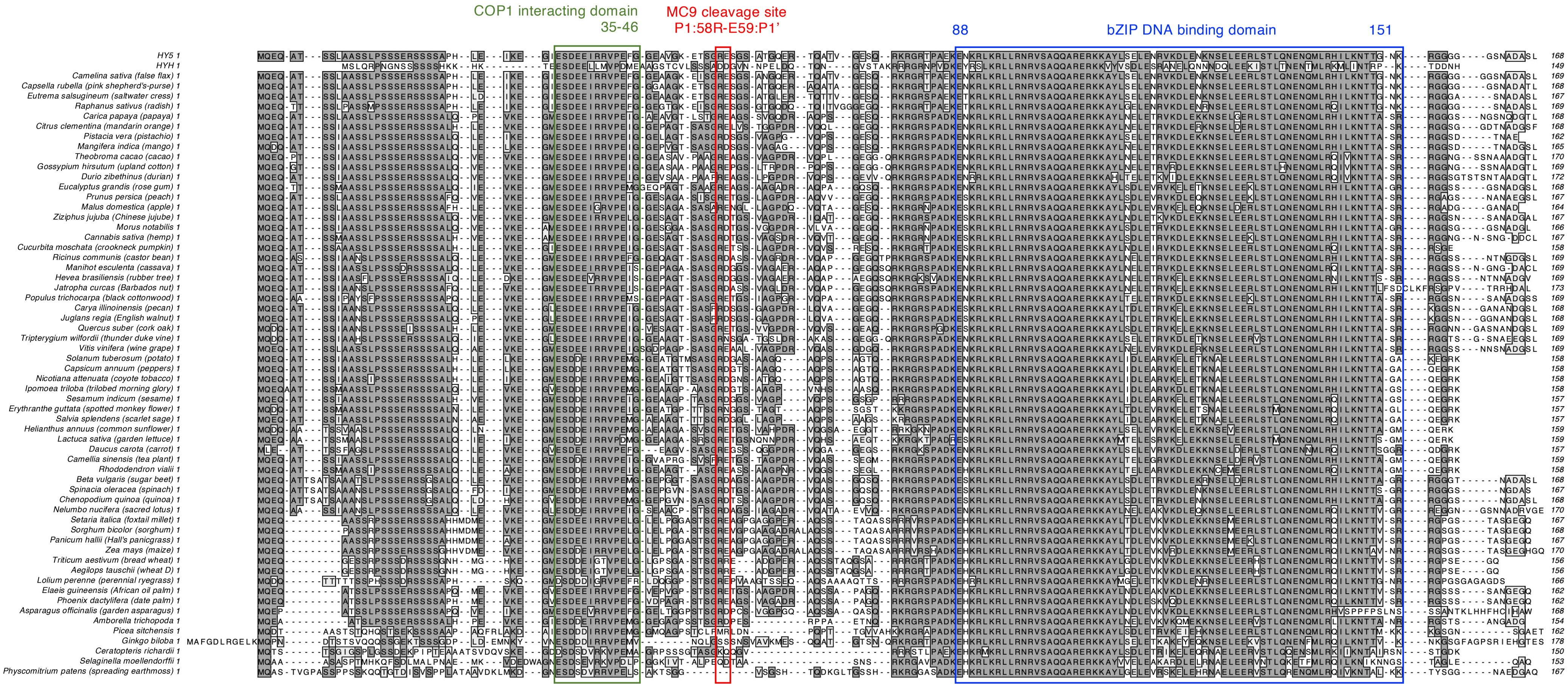

**Extended Data Figure 2: Diagrammatical representation of constructs used in this study.**
Protein domain in the Ubiquitin Fusion technique (UFT) construct are shown as coloured rectangles. Epitope tags are indicated in their relative positions in proteoforms. The effect and positions of residue substitutions are described. Promoter HY5 genomic region in constructs is drawn to scale.

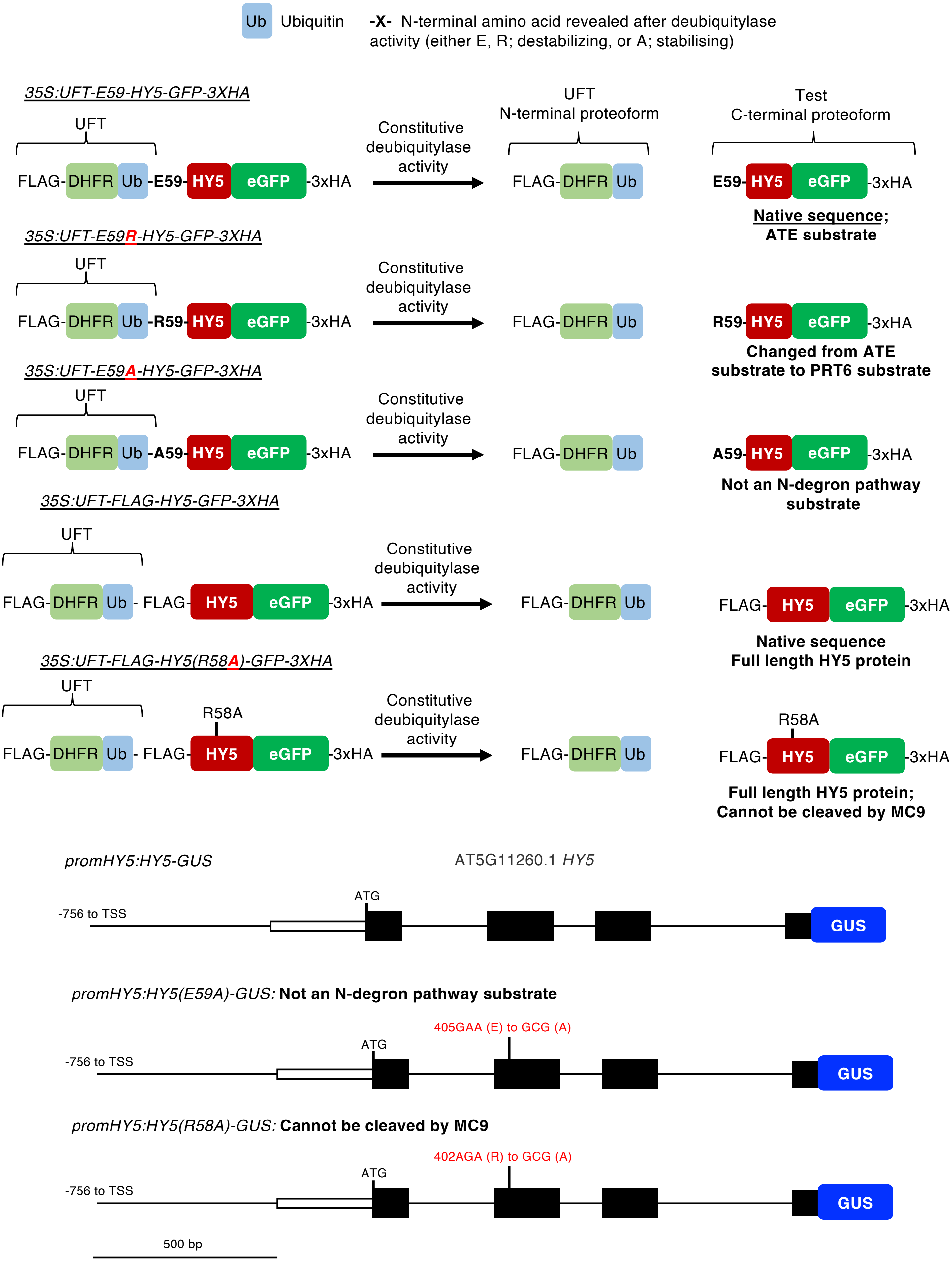

**Extended Data Figure 3:** Subcellular localization of E59-HY5-GFP-3xHA construct in different genetic backgrounds. E59-HY5-GFP-3xHA in *hy5-2* was crossed to other mutants to generate homozygous lines containing the same transgene integration event. Roots were assessed for GFP fluorescence in the presence or absence of bortezomib (BZ).

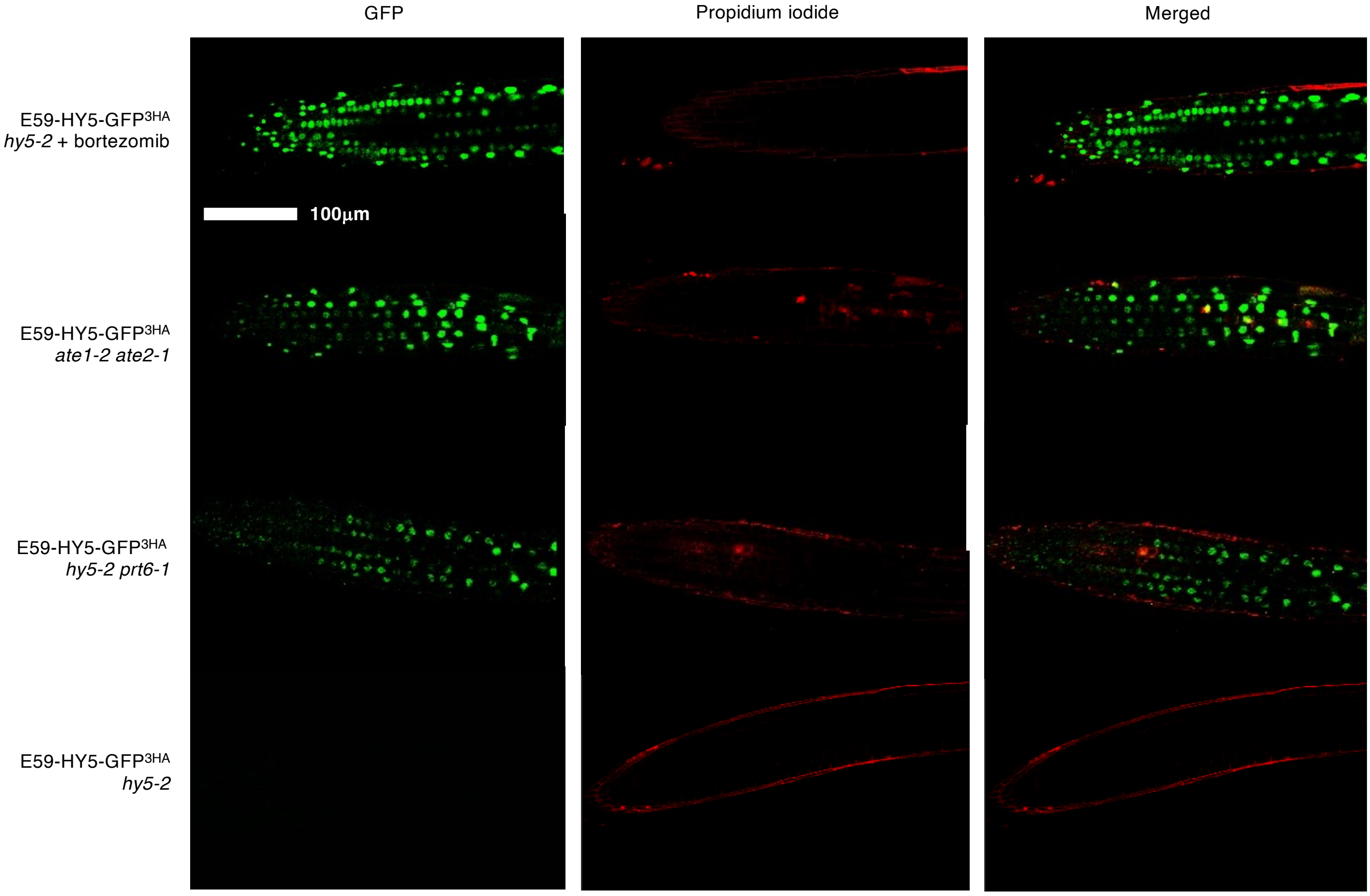

**Extended Data Figure 4:** Influence of *ATE1* and *ATE2* on *HY5* regulated hypocotyl elongation, and examples of phenotypes of seedling response to light.

**a.** Hypocotyl length of 7-day old seedlings grown in short days of WT, *hy5-2* and N-degron pathway mutant combinations. Data are presented as mean ± SD (n = 16) significant differences denoted with letters for one-way ANOVA (adjusted p<0.05).

**b.** Example images to show influence of *prt6-1* and E59-HY5-GFP-3HA on hypocotyl elongation under short days. 5-day old seedlings

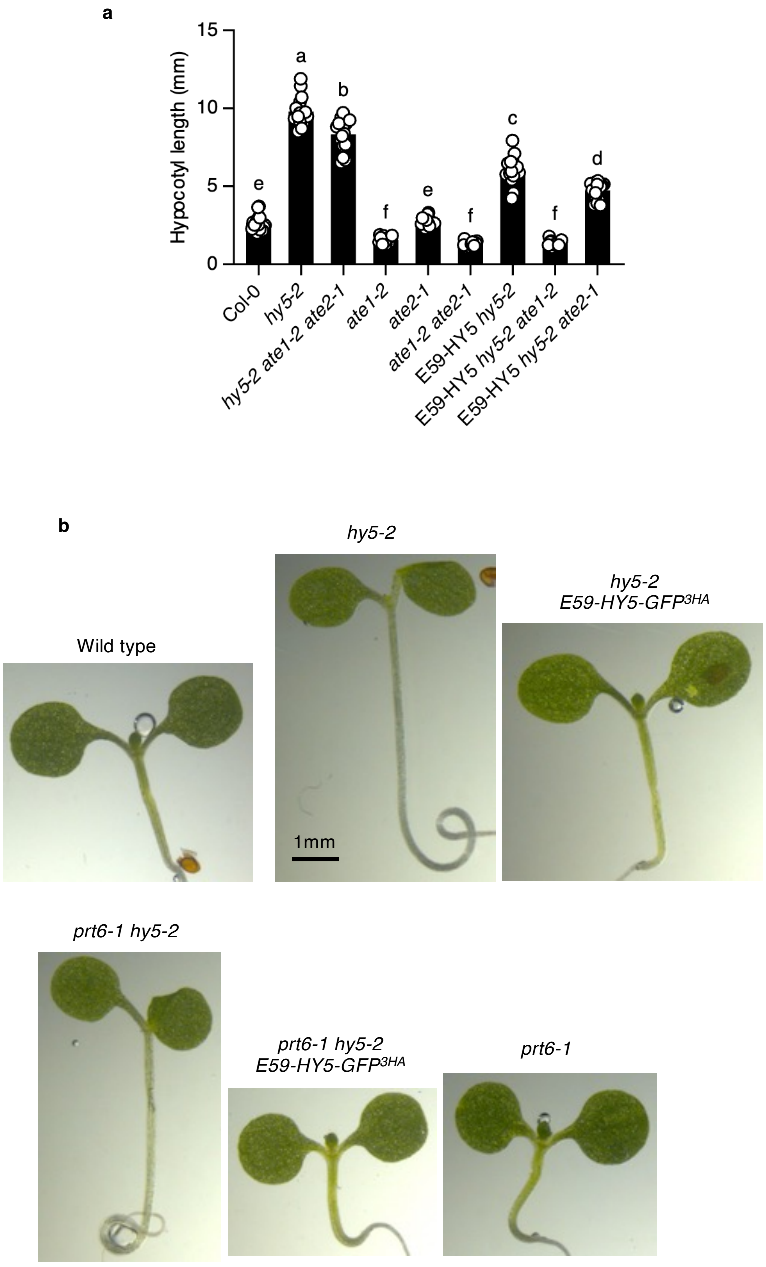

**Extended Data Figure 5:** Influence of N-degron pathway mutants on *HY5* regulated response to Tunicamycin.

**a.** Relative weights of 14-day old light grown seedlings of WT, *hy5-2* and N-degron pathway mutant combinations, seedlings grown in constant light on 60ng/ml Tunicamycin (TM). Data are presented as mean ± SD (n = 3) significant differences denoted with letters for one-way ANOVA (adjusted p<0.05).

**b.** Example images to show influence of *ate1-2* and *ate2-1* and *hy5-2* on seedling response to growth on Tunicamycin (60 ng/ml).

**c.** Example images to show influence of *prt6-1, hy5-2* and E59-HY5-GFP-3HA on seedling response to growth on Tunicamycin (60 ng/ml).

a

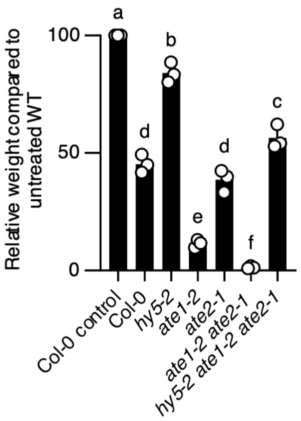


b

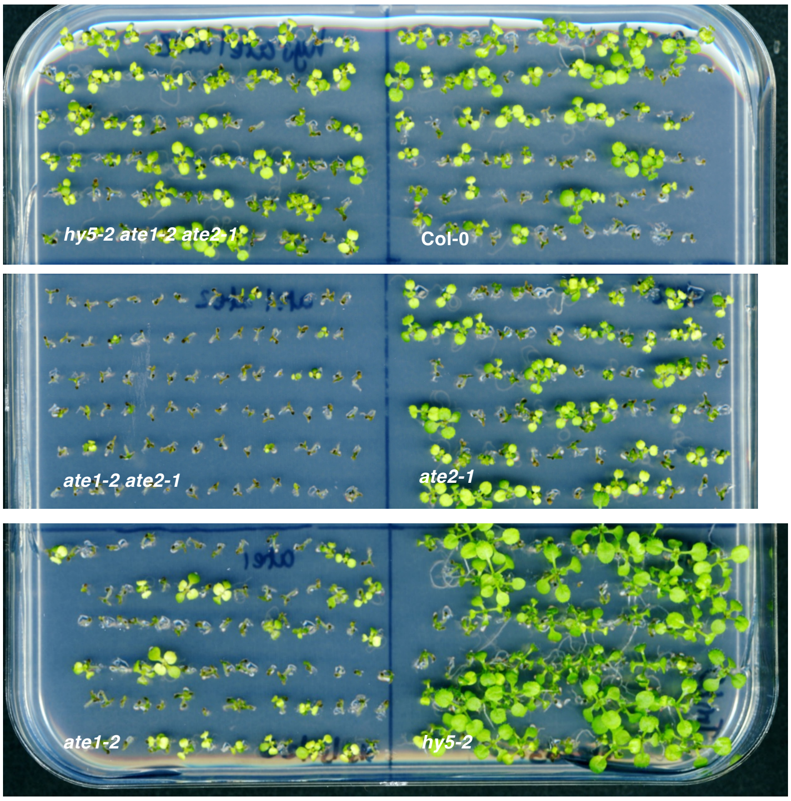


c

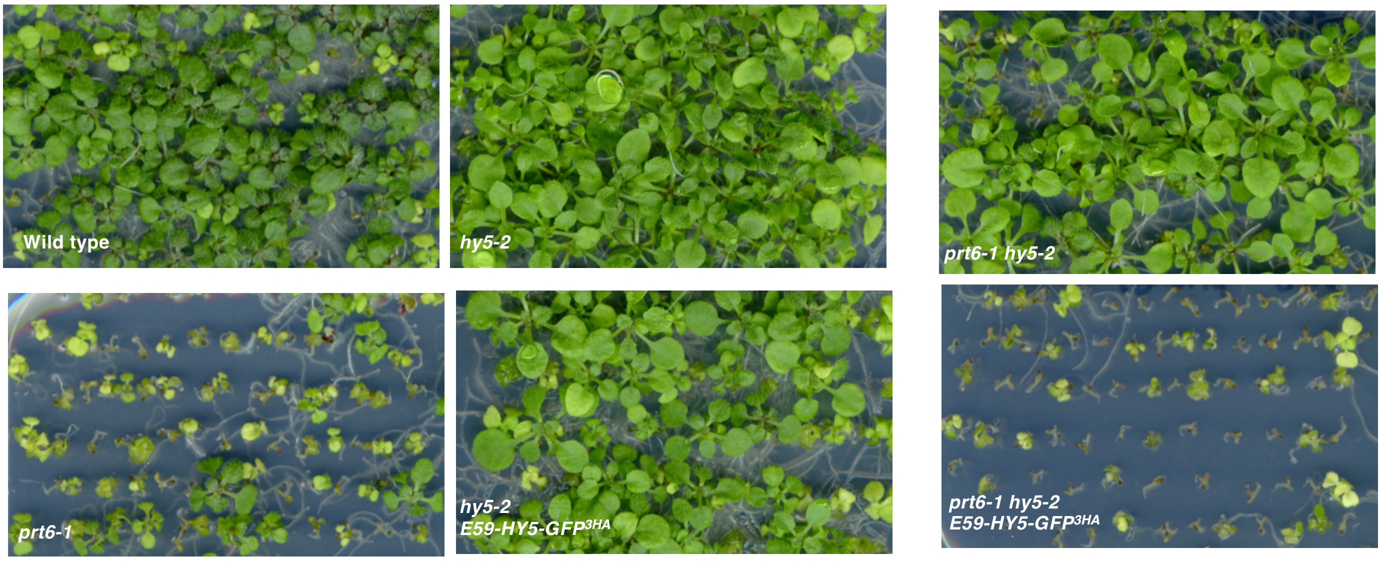

**Extended Data Figure 6:** Western blot analysis of HY5 and GUS constructs.
**a.** Stability of E59R-HY5-GFP-3HA in *hy5-2* in 5 day old etiolated seedlings following transfer from dark to constant light, or 5 days in constant light, in the absence or presence of bortezomib (BZ). Anti HA primary antibody.
**b.** Stability of E-, R-, or M-HA-GUS in 5 day old etiolated seedlings following transfer from dark to constant light in the absence or presence of bortezomib (BZ). Anti HA primary antibody.
**c.** Stability of promHY5:HY5 derivate proteins in 14-day old seedlings grown on Tunicamycin (TM; 60 ng/ml ) or control. Anti GUS primary antibody.
Ponceau staining of Western blot is shown (p). Open triangle, HA-tagged HY5 Ct-proteoform; closed square, full length HY5-GUS; open square, HY5-GUS C-terminal proteoform.

a

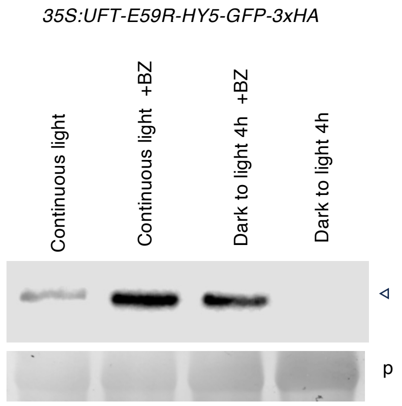


b

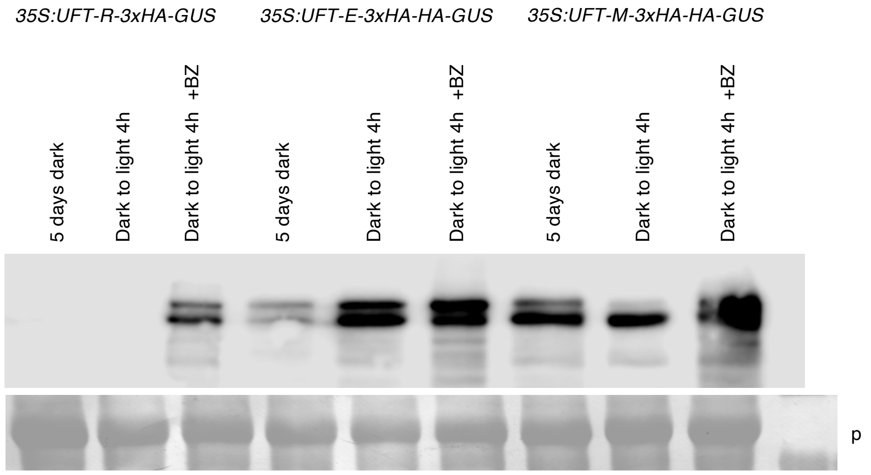


c

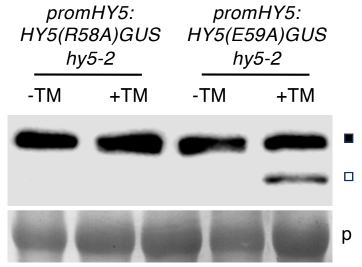

**Extended Data Figure 7 l** Expression of HY5 regulated genes in seedlings.
a. Expression of *XTH21* during the transition of 5-day old etiolated seedlings from dark to light. Data are presented as mean ± SD (n = 3).
b. Expression of *PDI* in 7-day old seedlings grown in constant light during liquid incubation in tunicamycin (5 μg/ml). Data are presented as mean ± SD (n = 3).

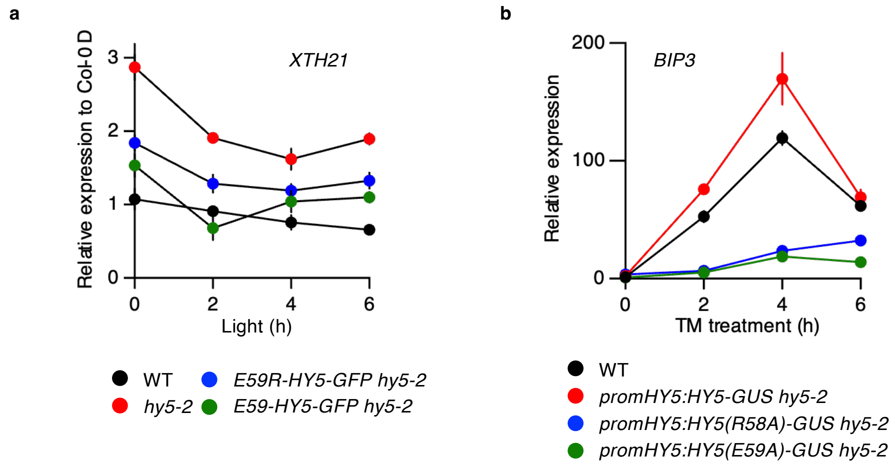

**Extended Data Figure 8:** Histochemical analysis of *promHY5:HY5-GUS* constructs in seedlings.
a. Seedlings grown in the absence of TM for 14 days
b. Seedlings grown in the presence of TM (60 ng/ml) for 14-days

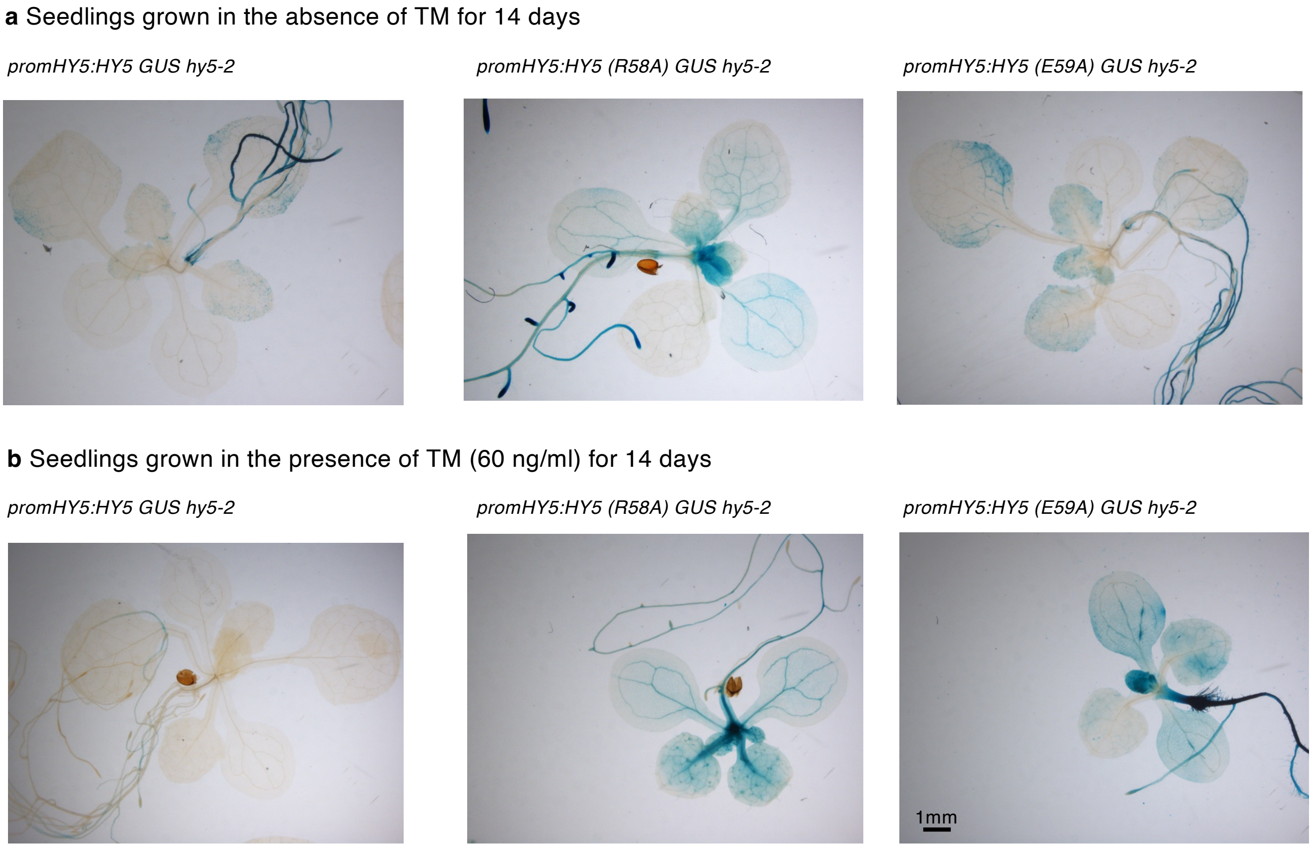
